# Vocal Transmission of Breeding Status May Facilitate Dispersal in a Cooperative Breeding Primate

**DOI:** 10.1101/525121

**Authors:** Robakis Efstathia, Watsa Mrinalini, Erkenswick Gideon

## Abstract

Social complexity may drive complexity in communicative systems due to an individual’s need to navigate unpredictable interactions with multiple conspecifics. Cooperative breeding primates (marmosets and tamarins; family: Callitrichidae) live in groups with moderate to high reproductive skew, particularly in females, whereby sexually mature individuals are frequently prevented from breeding. Remarkably, dispersal from natal groups is not stereotyped upon reaching reproductive maturity. Individuals are often observed remaining in their natal groups until the same-sex breeder in their group or a neighboring group dies, experiencing hormonal reproductive suppression, aggression, and limited access to potential mates. Here we examined whether emperor tamarins (*Saguinus imperator*) might use vocal signals to reduce dispersal risks and maximize the likelihood of attaining a breeding position. Using six consecutive years of mark-recapture data, we showed that sexually mature non-breeders (herein “secondary breeders”) are more likely to leave their groups from one year to the next than sexually mature breeders (“primary breeders”). This confirmed that, unlike primary breeders who do not need to disperse in order to reproduce, secondary breeders are choosing to accept the risks associated with dispersal and emigrating from their natal groups. We used neural networks to classify vocalizations according to individual breeding status, and conducted a series of playback experiments which demonstrated that tamarins discriminated between the calls of primary and secondary breeders. Our data support the hypotheses that secondary breeders disperse to increase mating opportunities and use vocalizations to signal their availability to potential mates. This species of cooperative breeder appears to use vocalization to navigate its social and reproductive systems, minimizing risks of dispersal and in turn increasing the likelihood of reproductive success. This research has important implications for our understanding of sexual signaling, partner choice, and reproductive success in cooperative breeders.

## INTRODUCTION

The social complexity hypothesis predicts that as social organization becomes more complex, so do systems of communication, in order to accommodate idiosyncratic interactions among conspecifics (Freeberg et al., 2012). Across taxa with differing levels of social complexity, there are consistencies in information transmitted in vocalizations such as individuality (e.g., elephants: Soltis et al., 2005; hyraxes: Koren et al., 2008; primates: Mitani et al., 1996; sea lions: Pitcher et al., 2012; wolves: Tooze et al., 1990) and sex (e.g., bats: Kazial & Masters, 2004; giant pandas: Charlton et al., 2009; marmots: Blumstein & Munos, 2005; primates: Rendall et al., 2004). However, vocalizations are also adapted to species’ own particular social pressures in ways that augment or replace social knowledge (Seyfarth et al., 2005). For instance, chacma baboons live in groups with strong dominance hierarchies and can distinguish the relative ranks of others based on their loud calls, which ostensibly lowers the risk of contact aggression (Kitchen et al., 2005). Similarly, geladas have developed a high number of vocalizations used specifically by males in affiliative interactions with the females in their harems (Gustison et al., 2012). Thus the communicative content of some vocalizations can reflect the unique needs imposed by social systems in which they are produced.

Cooperative breeding animals demonstrate a form of complex social organization that may drive the evolution of vocal complexity in order to accommodate the difficult task of group infant rearing (Hrdy, 2005; Zuberbuhler 2012; Leighton, 2017). Cooperative breeding systems are characterized by alloparental behavior—infant carrying and provisioning conducted by individuals other than the biological parents. Group social pressures are complex, since alloparents must reconcile their own fitness needs with the demands of the social group.

The Callitrichidae (marmosets and tamarins) are cooperative breeders who exhibit flexible social and reproductive strategies. Almost every type of social group has been observed within callitrichids: pair-bonded breeding couples, single females with multiple males, multiple females with multiple males, and, rarely, multiple females with a single male (Ferrari & Lopes Ferrari, 1989; Goldizen, 1996; Rylands, 1996; Watsa, 2015). However, most groups possess a single primary breeding female, and one or more breeding males (Garber, 1997; Watsa, 2015). Primary breeders are individuals who mate and can therefore potentially contribute to the next generation’s gene pool. In contrast, secondary breeders comprise those individuals in the group that are sexually mature non-breeders, who do not participate in a group’s mating system but provide alloparental care. Often, secondary breeders are the undispersed offspring of the social group (Nievergelt et al., 1999). The remainder of a group consists of non-breeders, or sexually immature offspring of the primary breeders (Koenig & Dickinson, 2016; Watsa et al., 2017).

Notably, there are no strict relationships between animal age, the onset of reproductive maturity, and dispersal behavior (Tardif, 1984). A secondary breeding tamarin may remain in its natal group, undergoing hormonal (Birnie et al., 2011; Castro & Sousa 2004; Saltzman, 2010) and/or behavioral (Mumme et al., 1983; Nelson-Flower et al., 2011; Young et al., 2006; Digby & Saltzman, 2009; Price & McGrew, 1991; Sousa et al., 2005) reproductive suppression, until it takes over as a primary breeder once a same-sex primary breeding adult has died (Lazaro-Perea et al., 2000; Yamamoto et al., 2014). Alternatively, it may use intergroup interactions to assess mating opportunities (Nichols et al., 2015; Caselli et al., 2018), or become a “floater” or “visitor” who has irregular or short-term associations with other groups, after which it may return to its natal group or continue to disperse permanently (Brown, 1969; Löttker et al., 2004; Watsa 2013). Dispersal, which is done by both sexes in callitrichids, can pose several risks, including increased susceptibility to predation and unfamiliarity with new territory (Goldizen, 1996; Ridley et al., 2008; Ronce, 2007).

In this context, the role of vocal signals in facilitating the dispersal of secondary breeders may be extremely important. Vocal sexual signaling has been explored in many taxa with various social systems, but our understanding of its role in cooperative breeding systems is incomplete (Andersson & Simmons 2006; Fitch & Hauser, 2003; Wachtmeister, 2001; Warrington et al., 2014). Callitrichids are highly vocal (Agamaite et al., 2015; Cleveland & Snowdon, 1982; Moody & Menzel, 1976; McLanahan & Green 1978; Masataka, 1982) and their calls contain information about the signaler such as age, sex, and individual identity (Miller et al., 2010; Norcross & Newman, 1993; Pistorio et al., 2006; Pola & Snowdon, 1975; Robakis et al., 2018). Long calls—high-amplitude long-distance vocalizations used for inter- and intragroup communication (Brown et al., 1979; de la Torre & Snowdon, 2009; Epple, 1968; Jorgensen & French, 1998; Lazaro-Perea, 2001; Ruiz-Miranda et al., 1999)—are good candidates for containing breeding status information because they can be received simultaneously by multiple individuals and groups. Unlike short-range signals, such as scent-marking, long calls can potentially facilitate partner and mate choice in a way that minimizes the dangers commonly associated with dispersal, including the likelihood of contact aggression with unfamiliar conspecifics (Crockford et al., 2007; McGregor, 2005).

In this study we collected mark-recapture, behavioral, and vocal data on a wild population of emperor tamarins (*Saguinus imperator*) to examine two hypotheses. First, we tested whether secondary breeders are less likely to remain in a group from year to year than primary breeders. Survival rates should be the same across all adult-sized individuals, and so elevated disappearances or group transfers in secondary breeders would indicate that individuals in that breeding class are emigrating; however, this potential disparity has not been explicitly tested in callitrichids. Second, we assess whether secondary breeders mediate the risks associated with dispersal by broadcasting and receiving breeding status information via their vocalizations. To test the first hypothesis, we used mark-recapture data from six consecutive years to test for differences in the rate at which primary and secondary breeders either disappear or disperse from a group from one year to the next. If secondary breeders are more likely to relocate than primary breeders, this would support the notion that secondary breeders do leave their groups more frequently to seek reproductive opportunities. To test the second hypothesis, we evaluated whether individuals differentiate between the vocalizations of primary and secondary breeders using playback experiments. We used only the long calls of unfamiliar female tamarins in this study because the influence of sex on reactions to unfamiliar intruders has not been consistent across studies (Lazaro-Perea, 2000; French et al., 1995; Caselli et al., 2018; French & Inglett, 1989). We also tested whether helper number influences reactions to the calls of unknown females. In a study of captive lion tamarins (*Leontopithecus rosalia*), the number of secondary breeders in a group was positively correlated with negative reactions to unfamiliar primary breeding females but had no effect on reactions to secondary breeding females (French & Inglett, 1989). This may be a strategy by which small groups discourage reproductive competition from primary breeding female immigrants while recruiting, or passively accepting, additional helpers to aid in infant rearing. This research is an important step toward understanding how communication evolves to accommodate complex social and reproductive systems with highly-contested breeding positions.

## METHODS

### Data Collection

Data collection took place at the Estación Biologica Rio Los Amigos (EBLA) in the Madre de Dios Department of Peru. Between 2014 and 2016 we recorded vocalizations from habituated groups of emperor tamarins (*Saguinus imperator*), who have been part of an ongoing mark-recapture and behavioral study since 2011 (Watsa et al., 2015). In short, tamarins were habituated to a multi-compartment trap over the course of 1-2 months before being captured as a group. If some individuals in the group were not habituated to the trap, the group was only captured without them if they were adults; infants and juveniles were never left alone while the rest of their group was being screened (Watsa et al., 2015). During animal processing, all individuals were given microchips to maintain identification across years; each tamarin was also fitted with a beaded collar that indicates group, sex, and identity, and given a unique bleach pattern on its tail, to facilitate identification during behavioral observation. Tamarin groups were released together, on the same day of capture, and well before sunset to ensure enough time in the day for foraging and locating a sleep tree. One female in each group, typically the dominant breeding female least likely to disperse or disappear, was given a radio collar (Wildlife Materials Inc.) to facilitate easy location of the group for subsequent behavioral observation and tracking.

### Determining Breeding Status

Breeding status was determined using a model based on morphological and behavioral observations (see Watsa et al., 2017 for a detailed description of the methodology). In brief, primary breeding females were initially identified by nipple lengths that indicated parity, and primary breeding males were identified according to whether researchers had noted copulations during behavioral observations. Secondary breeders were defined as undispersed individuals who had been born in the 1-2 years preceding capture and were nulliparous. Nonbreeders (juveniles and infants) were individuals born that year, identified by body size, deciduous dentition and facial pelage. During mark-recapture events, morphological measurements—including vulva length/width, testicular length/width, and body mass—were taken on all individuals. These measurements from individuals with known breeding statuses were used to train a model which then assigned breeding status to the rest of the individuals in the population based on variation in nipple lengths, body mass, vulvar index and testicular volume (Watsa, Erkenswick, & Robakis 2017).

### Relocations Across Breeding Status Classes

Using annual mark-recapture data from 2011-2017, we assessed the rate of relocations for individuals of each breeding status class. Marked tamarins who were not present in the population over consecutive years were designated as having “disappeared” regardless of the reason for their disappearance. An animal’s absence could be the result of dispersal to a group outside the population or death due to predation, poor health, or random events, but since we cannot be sure which occurred we assume that the rate of death is similar for all adult-sized individuals. Any significant difference in disappearance rate among breeding status classes could thus be attributed to dispersal. Individuals who remained in the trapped population but changed groups, and individuals who transferred into the population as adults, were counted as “dispersed” since their immigrations were confirmed. “Relocations” are therefore defined as the combined figure including both disappearances and dispersals. To accommodate low sample sizes, we used Fisher’s exact tests to determine if the proportion of disappearances and dispersals differed significantly between breeding status classes.

### Predicting Breeding Status from Long Calls Long call recording

Teams of two to three researchers collected data from 06:00 to 16:00 on two to five groups of emperor tamarins (totaling 19 individuals) each year from 2014-2016. They conducted 15-minute focal follows on all tagged individuals randomized to balance data collection across individuals and groups, resulting in 1,080 in-sight follow hours. Observer A recorded behavioral data collection into a small, handheld recorder (Sony ICD-PX312, Sony ICD-PX333, and Olympus VN-722PC models, mp3 format). To avoid the potential for human voices to obscure tamarin vocalizations, Observer B simultaneously recorded using a Zoom Handy Recorder with an accompanying shotgun microphones (Zoom H5 and H6 models, Zoom North America, Hauppage, NY) at the highest available sampling rate (44kHz/24-bit and 96-kHz/48-bit, respectively). When the focal animal vocalized, Observer A noted the tamarin’s behavior and confirmed the identity of the vocalizer. At the end of the vocalization, Observer B confirmed the identity of the producer into the Zoom recorder. Long calls of identified individuals were also recorded opportunistically during behavioral data collection. If an animal besides the focal individual vocalized during a focal follow, observers identified the producer and, if possible, its behavior at the time it produced the vocalization. If a focal individual was out of sight and another tamarin produced a long call, Observer B began an *ad libitum* focal on the vocalizing individual until the focal animal was found by Observer A, or until the *ad libitum* individual was out of sight, whichever occurred first.

### Analysis of Producer Characteristics in Long Calls

Spectrograms of all long calls were generated in Raven Pro using a Hann window, 5.33 ms window size, 1.2 ms hop size, and 2048 DFT (Bioacoustics Research Program, 2014). All spectrograms were initially visually inspected for quality: those with low signal-to-noise ratios, and those which included an acoustic disturbance that obscured the signal of interest (e.g. a sudden loud background noise) were removed from the sample.

For call measurement we followed the methodology of Robakis et al (2018). Thirty-four unique measurements were taken on each long call (Appendix 1), including 12 robust measurements automatically generated by Raven Pro. These measurements reduce variation introduced by user error by using signal-to-noise ratios as opposed to measurements taken manually by a researcher (Charif et al., 2010). Long calls are characterized by a series of disconnected syllables, and so we produced two sets of measurements, those on each syllable, or discrete subunit, within a call (“syllable set”), and those on the entire call (“unit set”). Each syllable set measurement was given a quality score from 0-3, with 0 indicating that none of the measurements were reliable (e.g. if the signal-to-noise ratio was too low at a given point in the syllable), and 3 indicating that all measurements were reliable. All unit set measurements were then given corresponding 0-3 scores. Only long calls that received a score of 3 for both syllable and unit sets were included in the sample.

We trained and ran artificial neural networks for each species based on all 34 measurements (Robakis et al., 2018). Artificial neural networks (ANNs) are machine learning algorithms that are trained using an iterative process to categorize inputs, such as frequency measures, as outputs, such as breeding status. The network is made of a hidden layer of “neurons”, or nodes, that get connected to inputs via weighted paths; the network, which runs completely without human supervision, assesses which configuration of inputs and weights-per-input results in a model with the best predictive accuracy. The final, most accurate model can then be used to classify novel inputs. Here we trained the network using one hidden layer, two to 10 neurons in increments of two (2, 4, 6, 8, 10), weight decays of 0.001, 0.01, 0.05, 0.1, and initial random weights of 0.5 and 1 (e1071 package: Meyer et al., 2012; nnet package: Venables & Ripley 2002) (R Core Team, 2017). To avoid overfitting the model, it was trained on a randomly selected 67% of the data (the training set) and run on the remaining 33% (the test set). Outcomes for ANNs are somewhat stochastic based on input data, so we ran the network on 10 unique test sets, each comprising a randomized subset of 33% of the original dataset, and used the mean of all results as the accuracy of the network (Robakis et al., 2018).

### Playback Experiments Stimulus preparation

Playback files were made using Audacity software (Audacity Team, 2015), and played on mp3 players (Tomameri mp3/mp4 player; GNBI Inc., Merrillville, IN) through Altec Lansing speakers (IMW576; New York, NY) via an auxiliary cable. Each file began with a three-second, 450-Hz pure tone to indicate the start of the file, followed by 20 seconds of silence to allow observers to distance themselves from the speaker. Starting at the 23-second mark, one-second 550-Hz tones played every minute for 10 minutes to indicate the start of scans. After the first 550-Hz tone, vocalizations played: experimental condition recordings consisted of three long calls separated by 10 seconds each, and the control vocalizations were a 17-second phrase of duetting titi monkeys (*Plecturocebus brunneus*). A three-second, 450-Hz tone signaled the end of the experiment. This was followed by another 20 seconds of silence, to allow the observers to locate the speaker, and a final three-second, 450-Hz tone to signal the end of the file. Each file was amplified to a maximum of −3 dB and exported as an mp3 file. Tamarins were not observed to attend to either the 450-Hz or 550-Hz tones across all experiments.

To avoid the potentially confounding effects of sex, we played long calls of primary or secondary breeding females to two habituated groups of emperor tamarins (Caselli et al., 2018; Lazaro-Perea, 2000). For primary breeding female (PBF) and secondary breeding female (SBF) conditions, we used long calls recorded from known individuals in 2015 and 2016, and the control recorded from a group of *P. brunneus* at EBLA in 2015. We chose the calls of *P. brunneus* because groups are often observed ranging near emperor tamarins (all authors, *pers. obs.*), and so the tamarins are regularly exposed to their vocalizations throughout the day. To reduce the likelihood that tamarins were reacting to the long calls of known conspecifics (Herbinger et al., 2009; Wich et al., 2002), we only used calls recorded from tamarins from non-neighboring groups. To avoid pseudoreplication, we created six files for each conspecific condition that could be played to each group (Kroodsma, 1986; Zuberbühler & Wittig, 2011). Each file presented long calls in a unique order, and all files were played to a group at least once before any were repeated. We ran experiments on groups SI-3 and SI-4, as we did not have long calls from both unfamiliar PBF and unfamiliar SBF individuals for groups SI-1, SI-2, or SI-5. Group SI-3 comprised four individuals: one PBF, one primary breeding male (PBM), one SBF who had been a juvenile in the previous year, and one untagged adult individual of unknown breeding status who was often seen with the group. Group SI-4 had nine individuals: one PBF, two PBMs, two secondary breeding males (SBMs) who had been juveniles the previous year, and two juvenile (nonbreeding) females who were not in the population the preceding year. Two untagged adult individuals of unknown breeding status were also regularly seen with that group.

### Experimental protocol

Playback experiments took place from July to August, 2017, between 06:00 and 16:00, on groups SI-3 and SI-4. Trials were only attempted if all the following criteria were met: no long calls were emitted by any emperor tamarins (either the focal group or a neighboring group) for a minimum of 20 minutes; observers had a minimum of two tagged individuals of any age class from the focal group in sight; and an intergroup interaction was not underway. No more than two trials were run on the same group per day, and trials were not run on the same group two days in a row. A second trial was not run on a group unless a minimum of one hour had elapsed since the first one, and the second stimulus was never the same category as the first stimulus presented that day, e.g. a PBF trial could only be followed by an SBF or a control. Trials were scheduled to maximize balance across groups and stimulus categories.

Experiments were conducted by teams of two to three researchers. So that subjects did not come to associate observers with vocalizations, one observer placed the speaker a minimum of 5 m away from the group and other researchers while the other observer(s) kept the group in sight. The researcher who placed the speaker used the initial 20-seconds of silence to move away from the speaker before the experiment began. Starting concurrently with the first long call, observers recorded one-minute scans every minute for 10 minutes, as marked by each 550-Hz tone. Behaviors were spoken into a handheld recorder and decoded into an Excel file at the end of the field season (ethogram, Appendix 2).

### Analysis of Playback Experiments

We collapsed all behaviors noted during data collection into a binary (“reactive” or “non-reactive”) response variable (Appendix 2). Visual scanning of the environment and attention toward the speaker were both regarded as reactive behaviors, as they are common callitrichid reactions to startling or potentially dangerous stimuli and are often used to assess reactions during playback experiments (Barros et al., 2004; Caine 1984; Kirchhof & Hammerschmidt 2004; Koenig 1998). Following earlier research, we also included scentmarking, and approaches toward or retreats away from the speaker as reactive behaviors (Masataka, 1987; Waser, 1977; Wich et al., 2002). Foraging, grooming, resting, and lateral, undirected movement neither toward nor away from the speaker were classified as non-reactive behaviors.

To test whether the proportion of reactive to non-reactive behaviors increased in response to experimental stimuli, we fit a generalized linear mixed model (GLMM) with binomial error and logistic link function using the lme4 package in R (Bates et al., 2015). As in other playback experiments (Kirchhof & Hammerschmidt, 2006; Windfelder, 2001), tamarin reactions were strongest immediately after stimulus presentation, and so we analyzed the first three minutes after stimulus presentation. We included experimental condition (control, PBF, SBF) and sex-breeding status class of the individual (PBF, PBM, SBF, SBM) as fixed effects, and individual identity (eight levels) as a random effect with random intercept. Neither untagged individuals nor nonbreeders were included in analyses. We did not implement stepwise inclusion of independent variables in the model, due to the increased chance of a Type I error (Mundry & Nunn, 2009), so a Wald Chi-Square was used to test the model for significance.

This study is part of an ongoing mark-recapture program that began at EBLA in 2009. Research permits were granted by the Directorate General of Forestry Services in the Ministry of Agriculture (SERFOR). Data collection follows the Association for the Study of Animal Behaviour’s Guidelines for the Use of Animals, the American Society of Primatologists’ Principles for the Ethical Treatment of Primates, and the American Society of Mammalogists’ Guidelines. Playback experiments were approved by the Institutional Animal Care and Use Committee of Washington University in St. Louis (Protocol No. 20160033); mark-recapture protocols were approved by the Institutional Animal Care and Use Committee of University of Missouri – St. Louis (Protocol Nos. 12-04-06 and 733363-5).

## RESULTS

### Relocation Rates Across Sex and Breeding Status Classes

Seventy-nine unique emperor tamarins were captured between 2011 and 2017. In total, there were 13 instances of relocations with known outcomes—individuals who either transferred groups within the population or appeared as adults from outside the population—and 40 instances of individuals disappearing with no known outcome.

Within cases of relocations with known outcomes, there were eight transfers by six individuals to other groups within the marked population. Only two individuals, one primary breeding female and one secondary breeding male, were found to have dispersed twice within the marked population; each of these emigrations was counted as a single instance, totaling four dispersals for these two individuals. Of the eight transfers, four were done by secondary breeders and four were done by primary breeders. Two of the primary breeding dispersers, one male and one female, were primary breeders during the first year of the mark-recapture program (2011), and so we cannot know whether their dispersals were emigrations from their natal groups. Five individuals—one primary breeder (male) and four secondary breeders (two male and two female)—transferred into the marked population as adults. These individuals did not disperse within the trapped population again. Of 13 confirmed dispersals, 62% (eight) were secondary breeders, and 38% (five) were primary breeders.

There was a significant difference between the rate of relocations in male and female non-breeders (Fisher’s exact test, P < 0.005), but not within primary or secondary breeders (Table 1, Table 2). All non-breeding individuals qualified as having disappeared, as none were confirmed as having dispersed to another group within the marked population. Pooled-sex secondary breeders relocated significantly more than pooled-sex primary breeders: mean ± SD relocations per year for secondary breeders: 3 ± 1.97, N = 20; mean ± SD relocations per year for primary breeders: 2 ± 3.33, N = 15 (Fisher’s exact test: *P* = 0.001).

**Table 1.** Number of relocations and total number of emperor tamarin individuals trapped between 2010 and 2017.

|  | *Relocated* | *Remained* | *Ratio* | *Pooled Ratio\** |
| --- | --- | --- | --- | --- |
| **PBF** | 5 | 21 | 0.24 | 0.26 |
| **PBM** | 11 | 41 | 0.24 |  |
| **SBF** | 12 | 19 | 0.53 | 0.67 |
| **SBM** | 12 | 17 | 0.56 |  |
| **NBF** | 3 | 19 | 0.16 | 0.52 |
| **NBM** | 10 | 6 | 1.67 |  |
| **Total** | 53 | 123 | 0.43 |  |

\* Ratio of individuals who relocated to those who remained in the marked population. PBF = primary breeding female; PBM = primary breeding male; SBF = secondary breeding female; SBM = secondary breeding male; NBF = non-breeding female; NBM = non-breeding male.

**Table 2.** Results of Fisher’s exact tests for proportion of emperor tamarins who relocated from or into the tagged population from 2011 to 2017 based on sex and breeding status.

| **Sex and Breeding Status** | **P-value** |
| --- | --- |
| NBF vs NBM† | **< 0.005** |
| SBF vs SBM† | 1.000 |
| PBF vs PBM† | 1.000 |
| (PBF + PBM) vs (SBF + SBM) † | **0.015** |
| (SBF + SBM)† vs (NBF + NBM) | 0.670 |
| (PBF + PBM) vs (NBF + NBM) † | 0.117 |

Significant results are bolded, and † indicates which sex and breeding status group had higher relocation rates. PBF = primary breeding female; PBM = primary breeding male; SBF = secondary breeding female; SBM = secondary breeding male; NBF = non-breeding female; NBM = non-breeding male.

### Predicting Breeding Status from Long Calls

From 2014-2016 we collected 205 long calls from 19 individually identifiable emperor tamarins from five groups (Table 3). Neural network accuracy based on the classification subset for categorizing long calls according to breeding status was 90.3% for the syllable set and 75.9% for the unit set.

**Table 3.** Number of individuals per breeding status class in the study populations of emperor tamarins whose long calls we recorded from 2014-2016.

| Year | Group | PBF | PBM | SBF | SBM | NBF | NBM | N |
| --- | --- | --- | --- | --- | --- | --- | --- | --- |
| **2014** | SI-3 | - | - | - | - | - | 1 (2) | 2 |
|  | SI-4 | 1 (2) | - | - | - | - | - | 2 |
| **2015** | SI-1 | 1 (10) | 2 (13) | 1 (5) | - | - | 1 (15) | 43 |
|  | SI-2 | - | 1 (16) | 1 (8) | 1 (5) | - | - | 29 |
|  | SI-3 | - | - | - | 1 (2) | - | - | 2 |
|  | SI-4 | 1 (7) | - | - | - | - | - | 7 |
|  | SI-5 | 1 (3) | - | - | - | 1 (1) | - | 4 |
| **2016** | SI-1 | 1 (16) | 1 (1) | 1 (12) | - | - | - | 29 |
|  | SI-2 | 1 (9) | 1 (13) | 2 (16) | 1 (1) | - | - | 39 |
|  | SI-3 | 1 (9) | - | - | 1 (7) | - | - | 16 |
|  | SI-4 | 1 (13) | - | - | 1 (15) | - | 1 (4) | 32 |
| **2014-2016** | **N** | **69** | **43** | **41** | **30** | **1** | **21** | **205** |

Number of vocalizations analyzed for each age-sex class presented in parentheses. PBF = primary breeding female, PBM = primary breeding male, SBF = secondary breeding female, SBM = secondary breeding male, NBF = nonbreeding female, NBM = nonbreeding male.

### Playback Experiments

We completed a total of 23 playback experiments: seven PBF conditions, 10 SBF conditions, and six control conditions. There were no aggressive or scentmarking behaviors observed, and so reactive behaviors comprised scanning, attention, approach, and withdraw (Appendix 2). Both experimental conditions elicited higher rates of reactive responses than the control condition (Table 4). Rates of reactive behaviors in response to SBF calls were higher than those in response to PBF stimuli (Wald Chi-Square: χ^2^ = 22.2, df = 2, *P* < 0.001, N = 135) (Figure 1, Table 5).

**Table 4.** Proportion of total behaviors exhibited by emperor tamarins following playback stimuli that were classified as reactive or non-reactive (ethogram: Appendix 2).

| Group | Reaction | Condition |  |  |
| --- | --- | --- | --- | --- |
|  |  | Control<br>(N = 6) | PBF<br>(N = 7) | SBF<br>(N = 10) |
| SI-3 | Reactive | 0.20 | 0.43 | 0.63 |
|  | Non-Reactive | 0.80 | 0.57 | 0.38 |
| SI-4 | Reactive | 0.17 | 0.23 | 0.75 |
|  | Non-Reactive | 0.83 | 0.77 | 0.25 |
| Combined Groups | Reactive | 0.18 | 0.36 | 0.67 |
|  | Non-Reactive | 0.82 | 0.71 | 0.33 |

Control = vocalizations from sympatric titi monkeys (*Plecturocebus brunneus*); PBF = long calls of primary breeding female emperor tamarins; SBF = long calls of secondary breeding female emperor tamarins.

**Table 5.**
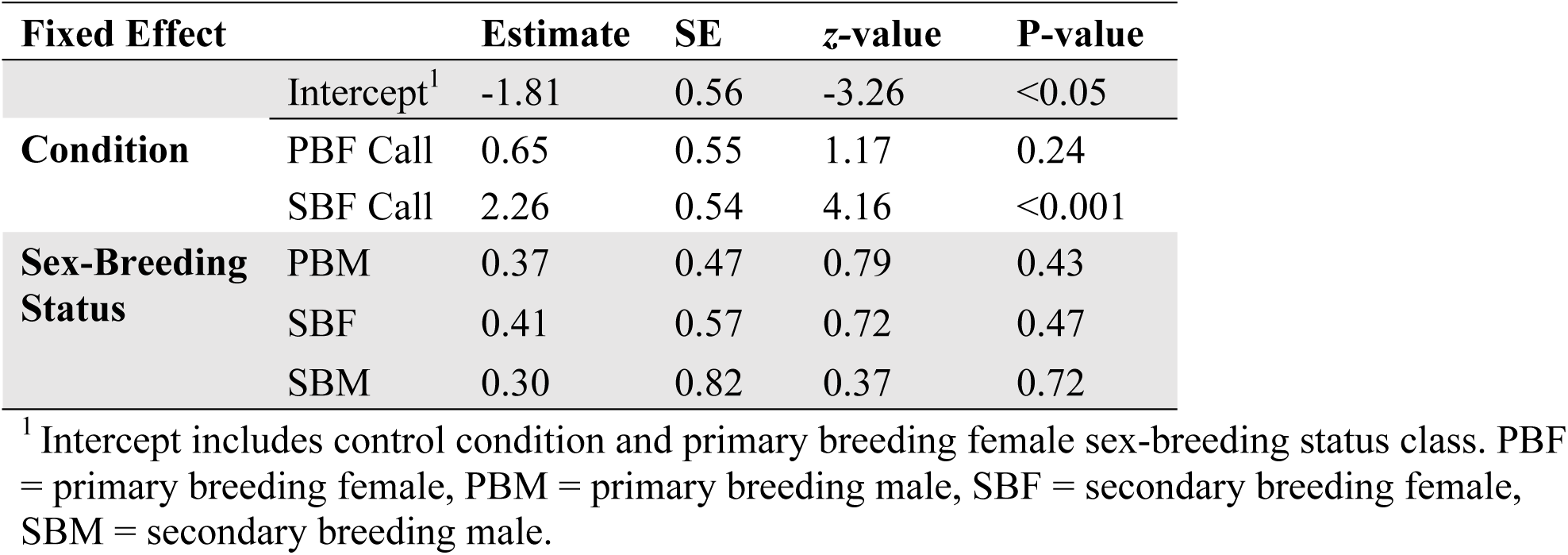
Influence of fixed effects on behavioral responses by emperor tamarins to playback stimuli.

| Fixed Effect |  | Estimate | SE | z-value | P-value |
| --- | --- | --- | --- | --- | --- |
|  | Intercept<sup>1</sup> | -1.81 | 0.56 | -3.26 | <0.05 |
| **Condition** | PBF Call | 0.65 | 0.55 | 1.17 | 0.24 |
|  | SBF Call | 2.26 | 0.54 | 4.16 | <0.001 |
| **Sex-Breeding Status** | PBM | 0.37 | 0.47 | 0.79 | 0.43 |
|  | SBF | 0.41 | 0.57 | 0.72 | 0.47 |
|  | SBM | 0.30 | 0.82 | 0.37 | 0.72 |

<sup>1</sup> Intercept includes control condition and primary breeding female sex-breeding status class. PBF = primary breeding female, PBM = primary breeding male, SBF = secondary breeding female, SBM = secondary breeding male.

**Figure 1.**
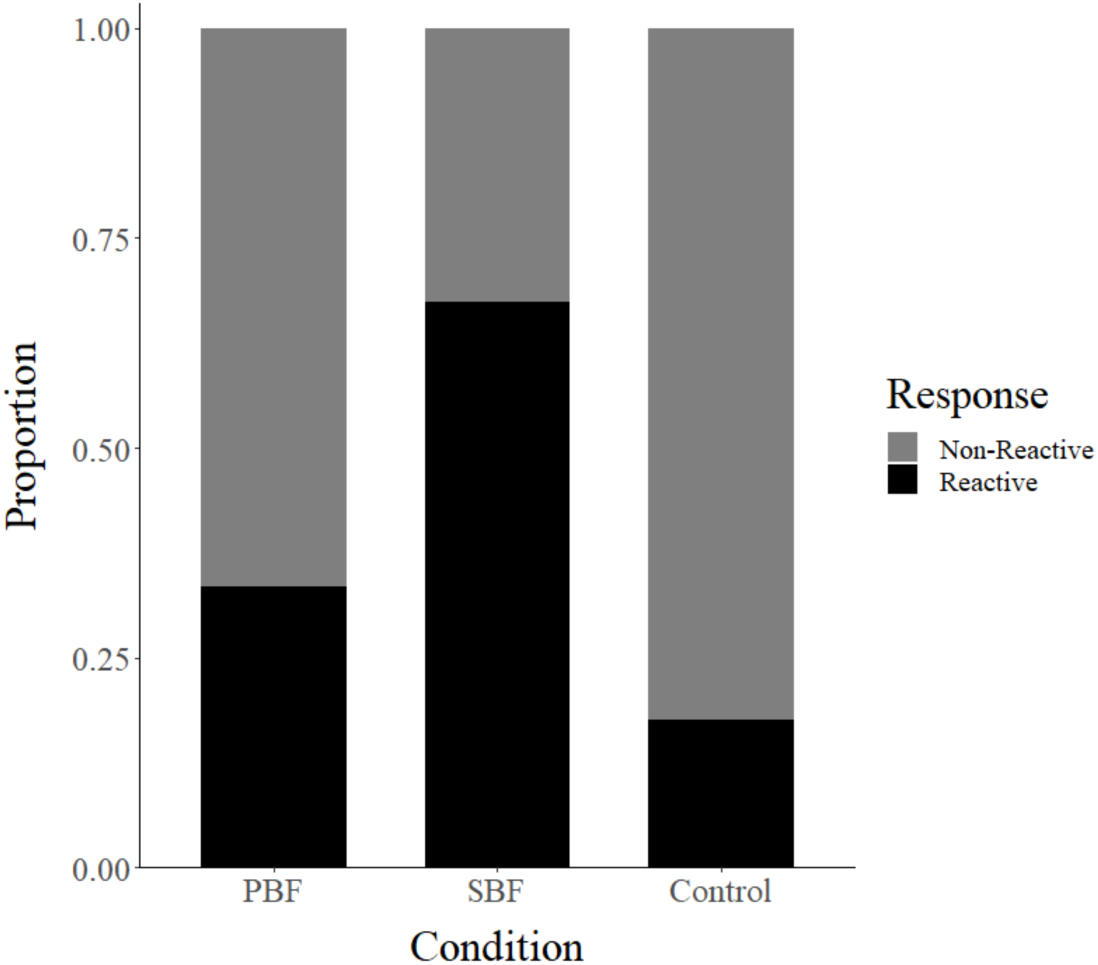
Proportion of total behaviors categorized as reactive or non-reactive (see Appendix 1 for definitions) exhibited by emperor tamarins in response to the vocalizations of primary breeding female emperor tamarins (PBF), secondary breeding female emperor tamarins (SBF), and the vocalizations of sympatric titi monkeys (*Plecturocebus brunneus*, control).

We were not able to detect an effect of individual identity on behavior (variance = 0 ± 0 SD). When pooled, females in this study were not more likely than males to react more to the vocalizations of the calls of unfamiliar females: 45% of female and 52% of male responses were reactive (Fisher’s exact test: *P* = 0.55). When compared to primary breeding females, PBMs and SBFs were slightly more reactive, and SBMs were slightly less reactive, but the effect of breeding status on reactive behaviors was not significant (Wald Chi-Square: χ^2^ = 0.77, df = 3, *P* = 0.86, N = 135) (Table 5). When only primary and secondary breeding playback conditions are compared, however, secondary breeders were more reactive than primary breeders to the PBF condition (Figure 3, Appendix 3). The proportion of reactive behaviors by secondary breeders was lower than that of primary breeders in response to the SBF condition. We predicted that Group SI-3 would exhibit a lower proportion of reactive behaviors in response to primary breeding females than SI-4, which had more helpers, but the opposite was true; group SI-4, however, was slightly more reactive toward SBF long calls than to PBF calls, though not significantly so (Figure 2, Table 4).

**Figure 2.**
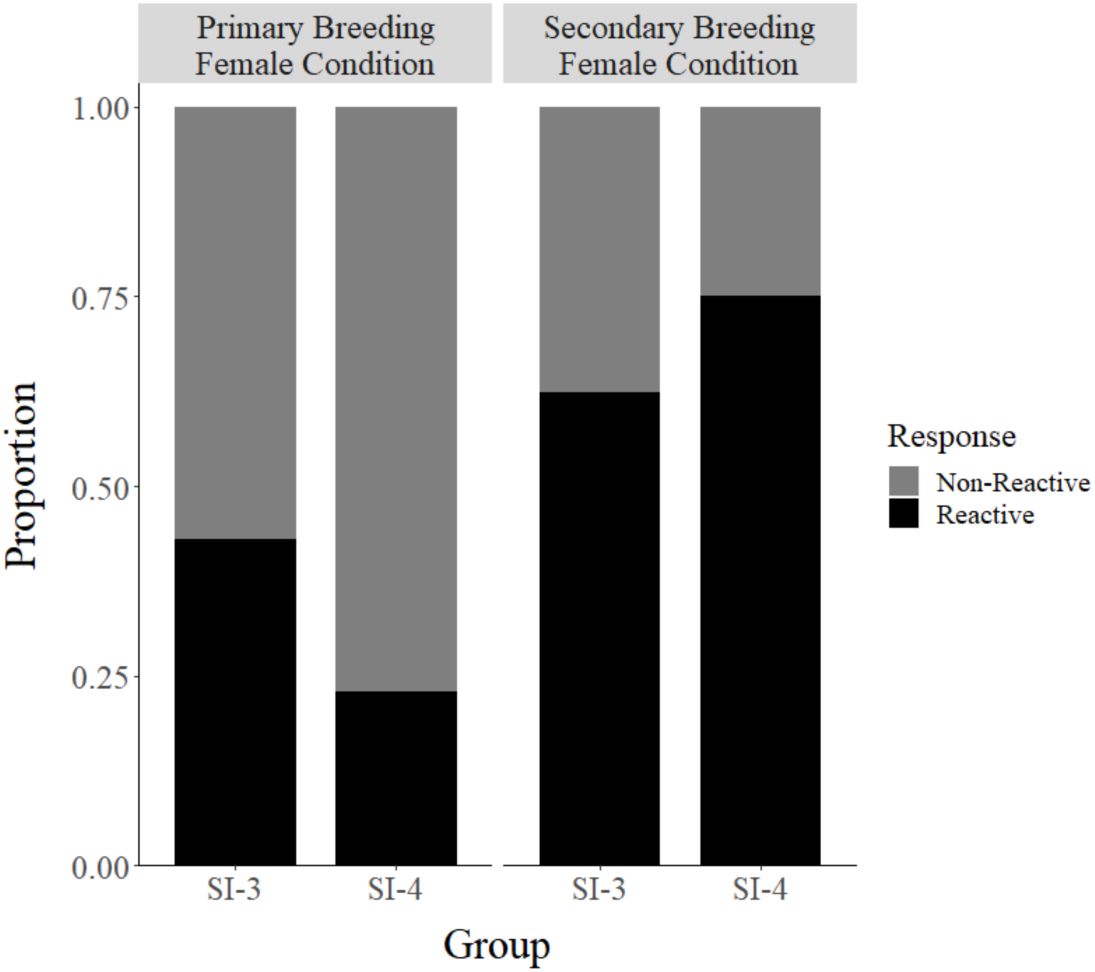
Proportion of reactive and non-reactive behaviors (see Appendix 2 for definitions) exhibited by two groups of emperor tamarins in response to primary breeding female and secondary breeding female long calls. Group SI-3 included a breeding pair, one secondary breeder, and one juvenile; group SI-4 included three primary breeders, two secondary breeders, and two juveniles. We predicted that groups with larger numbers of adult helpers would exhibit a higher proportion of reactive behaviors than groups with few helpers, and our predictions were not met.

**Figure 3.**
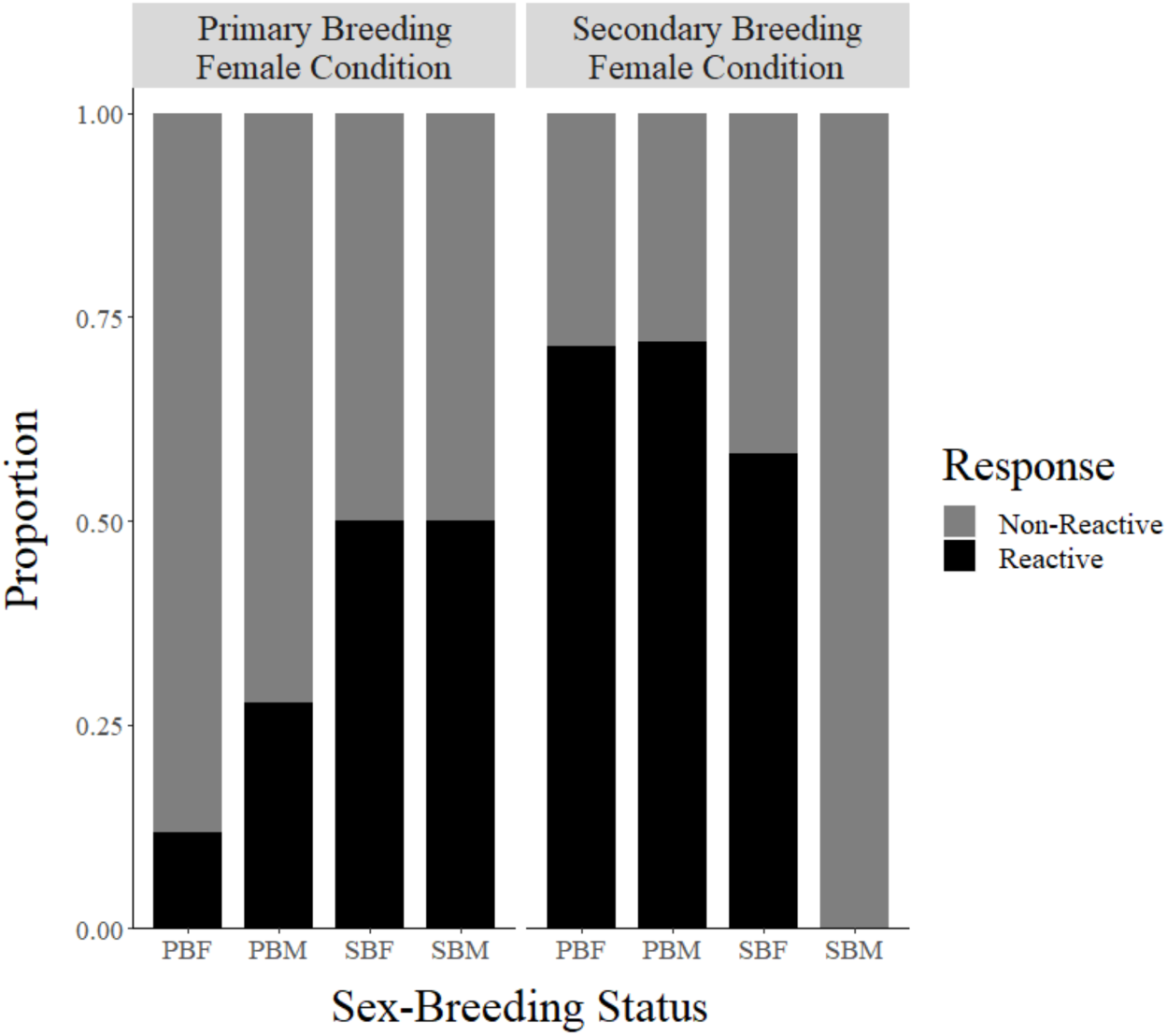
Proportion of reactive and non-reactive behaviors (see Appendix 2 for definitions) exhibited by emperor tamarins of each sex-breeding status class in response to playback experiments. PBF = primary breeding female, PBM = primary breeding male, SBF = secondary breeding female, SBM = secondary breeding male.

## DISCUSSION

The results presented here support the idea that vocalizations may play a role in facilitating emperor tamarin dispersal. Our expectation that secondary breeders are more likely to relocate from one year to the next than primary breeders was supported, most likely reflecting that they are dispersing. Our results demonstrated no statistical difference between relocation rates of secondary breeding males and females. This confirms that, after accounting for deaths, which should affect both sexes equally, males and females disperse at roughly equal rates (Goldizen, 1996). We found that the relocation rate for non-breeders exceeded that of primary breeders, likely due to infant mortality (Watsa et al., 2017). However, within non-breeders, males relocated at significantly higher rates than females, which may suggest that males disperse earlier than females, and earlier than previously reported in other callitrichids (Löttker et al., 2004; Goldizen & Terborgh, 1989). We confirmed that long-distance vocalizations can be accurately discriminated by breeding status and that individuals were significantly more reactive to the calls of secondary breeding females than primary breeding females. However, contrary to our prediction, we found no evidence that higher helper number is positively correlated with reactivity rates. The higher rates of reactive behaviors from group SI-3 towards PBF stimuli might be a generalized response of a small group to an unfamiliar individual in their territory (Peres, 1989). Although SI-4 did exhibit more reactive behaviors to SBFs than SI-3, the difference is much smaller than the difference between the groups’ reactions to PBF calls and may disappear with a larger sample.

There are several reasons for secondary breeders to immigrate from their natal groups. First, the role of morphological development cannot be discounted in younger males, whom we found to relocate significantly more than females. Löttker et al. (2004) reported a minimum of 1.4 years of age (maximum ≥ 3.7 years; average 2.4 years) for natal dispersers in mustached tamarins (*Saguinus mystax*). In a longitudinal study of saddleback tamarins (*Leontocebus weddelli*) Goldizen and Terborgh (1989) found that the youngest dispersers of confirmed age were 1.3-1.5 years old, but also noted that the youngest immigrant was male. In keeping with these findings, the model developed by Watsa et al. (2017) to predict breeding status did not include individuals under one year old as potential breeders based on their young age. Earlier research on this population demonstrated that male emperor tamarins develop scent glands, which may play a role in sexual behaviors (French et al., 1984; Heymann 2001; Miller et al., 2003), at about 6 moths of age, while females develop theirs around 1.5 years (Watsa, 2013). Moreover, though there was not comprehensive data on testicular volume for this population of emperor tamarins, the same earlier study included sympatric saddleback tamarins (*Leontocebus weddelli*), which demonstrated that by 1.5 years of age testicular morphology appears adult while vulvar morphology does not (Watsa, 2013). Emperor tamarins generally give birth between November and March (Watsa, 2013); as such, the oldest individuals in a cohort trapped between June and August would be 7-9 months of age—old enough for males, unlike females, to have highly developed scent gland and testicular morphology while remaining under the one-year threshold. This difference in development may stimulate earlier dispersal in males than females, causing a sex difference in relocations in the non-breeder class that then disappears by 1.5 years of age when both sexes are morphologically adult.

Male and female secondary breeding callitrichids are also highly susceptible to marginalization and aggression from same-sex adults (French & Inglett, 1989; Sousa et al., 2005; Ginther et al., 2001). In the emperor tamarin population at EBLA, we have observed behaviors and features of certain individuals that are consistent with expectations for secondary breeders in the process of dispersing. The majority of trapped individuals in our mark-recapture program are re-trapped the subsequent year if they are still in the population (Watsa et al., 2015), and those few who are not re-trapped share a suite of characteristics. They are adult-sized, and habituated to trap sites, suggesting that they did not immigrate from outside of the trapped population. They tend to associate with a single group but keep a distance, and forage and sometimes even sleep several meters away from the rest of the group; they also typically wait to visit the trap site to feed after the rest of the group has already done so (all authors, *pers. obs.*). These avoidant behaviors appear to be a strategy by which individuals avoid aggression from group members. Our observations strongly suggest that they are secondary breeders dispersing from their natal groups. In addition to experiencing marginalization and aggression, waiting to occupy the position of a primary breeder in their natal group who has died exposes offspring to potential inbreeding (Cooney & Bennett, 2000; Riehl, 2017). Reports of copulations of adult females with their male offspring in the wild are exceedingly rare (Goldizen, 1996), and one study of golden lion tamarins revealed that no infants that resulted from a female mating with her father or brother survived to weaning (Dietz & Baker, 1993).

Under these circumstances, emigration would be one strategy by which callitrichids could increase their chances of reproducing. However, dispersal can be costly due to increased risks associated with high predation rates and unfamiliarity with new territory. Immigration to another group or the formation of a new group also both require acceptance by other individuals; in this sense, successful dispersal is inherently cooperative (Noë & Hammerstein, 1994). Zuberbühler (2016) has suggested that immigrating female chimpanzees have developed vocal strategies to reduce the likelihood of aggression from females. Secondary breeding callitrichids may similarly use vocal strategies to reduce the risks inherent to dispersal. For example, tamarins may not recognize an individual who has come from a non-neighboring group: detecting its breeding status from its vocalizations, especially when combined with its sex, might inform a group or individual’s receptivity to the immigration. Long-range signals would mitigate the risk of contact aggression that would otherwise be incurred during the exchange of short-range signals such as scent-marking. The complexities and flexibility of the cooperative breeding system renders communication between potential partners, or mates, critical to reproductive success, and offers a mechanism by which communicative complexity can be transmitted: individuals who “cooperate” vocally might be more likely to successfully reproduce.

Our current understanding of the role of vocalizations in cooperative breeding social groups is limited to one study on apostlebirds in which the authors suggest that transmitting breeding status in vocalizations uttered near the nest may facilitate mate recognition and alloparenting behaviors (Warrington et al., 2014). Research on the relationship between breeding status and vocalizations in marmosets and tamarins has focused on infants, who emit vocalizations to solicit attention from a parent or alloparent (Epple, 1968; Bezerra & Souto, 2008). Yet the role of vocal signals in facilitating partner recognition and mate choice in adults have been explored in many taxa with various social systems besides cooperative breeding (Andersson & Simmons 2006; Fitch & Hauser, 2003; Wachtmeister, 2001). To date, studies on primate vocalizations and mate choice have tended to focus on intragroup signals in large groups (e.g. copulation calls: Semple, 2001), interindividual signals in solitary species (e.g. estrus calls: Buesching, Heistermann, Hodges, & Zimmermann, 1998), or on mate defense and the strengthening of pair-bonds (Caselli et al., 2014; Cowlishaw, 1996; Geissmann & Orgeldinger, 2000). Historically, sexual signals have been defined in part by their sexual dimorphism (Darwin, 1871; Andersson, 1982).

Although callitrichine long calls are not canonically dimorphic, ornamented sexual signals, nor are they overtly different to the human ear, they do differ significantly by sex (Masataka, 1987; Norcross & Newman, 1993; Robakis et al., 2018). Given that long calls can and do serve multiple socially important functions in primates, such as intragroup coordination and resource defense (Wich & Nunn, 2002), it is possible that callitrichine long calls act as sexual signals in addition to other performing other functions (Bergman & Sheehan, 2013; Casselli et al., 2018). In a population where reproductive success is limited by both the social and mating systems, it would benefit potential dispersers to transmit and receive information about breeding status. An expansion of our assumptions of how sexual signals are used in primates can lead to a clearer understanding of the role of vocalizations in dispersal and mate choice.

Ultimately, these results are a first step toward understanding proximate behaviors tied to cooperative breeding and dispersal in callitrichids. Future studies should be undertaken to increase sample sizes to clarify whether the presented pattern of sex-breeding status effects on reactions holds, and should consider whether experiments are conducted in the core or periphery of a group’s home range (Caselli et al., 2018). They should also focus on playing the long calls of opposite-sex individuals of both breeding statuses to secondary breeders alone. This will help elucidate whether dispersers are truly more likely to attend to calls of individuals they may regard as potential mates. Another important next step in understanding the role vocalizations in partner and mate choice in cooperative breeders must involve experiments to examine the intentionality behind the production of long calls by secondary breeders. Based on the nonlinear relationship between age and breeding status (Watsa et al., 2017) and on a prior study of the same dataset which accurately classified calls according to age-class (Robakis, Watsa, & Erkenswick, 2018), growth and development alone are unlikely to be the sole drivers of changes in vocalizations. Additional research will clarify exactly how vocalizations might be used during dispersal and confirm whether the relationship between spectrotemporal changes and dispersal behaviors are causal. That vocal parameters are not necessarily correlated with age demonstrates that something besides growth or senescence is driving these changes, but they may be hormonal rather than consciously produced. Research on food calls (Caine et al., 1995; Roush & Snowdon, 2000), antiphonal calling (Toarmino et al., 2017) and cooperation (Cronin et al., 2005) has shown that callitrichids do alter their vocalizations based on their perception of others’ knowledge and intent. Research targeting the development of vocalizations of secondary breeders would clarify the bases of these changes, and the use of these vocalizations as sexual signals for both males and females.

## ACKNOWLEDGEMENTS

We thank the Peruvian Ministry of Agriculture, the staff at EBLA, and the Amazon Conservation Association for making this work possible. We are also grateful to each of the research assistants involved in data collection. This research was funded by the American Society of Primatologists, the International Primatological Society, Field Projects International, and Washington University in St. Louis.

### Appendix 1

Syllable and unit parameters measured from long calls produced by emperor tamarins. Parameters marked by † are robust signal measurements automatically generated by Raven Pro. Abbreviations for each variable are in parentheses. Adapted from Robakis et al., 2018.

| **Syllable Set** |  |
| --- | --- |
| ***Variable*** | ***Description*** |
| Age Class | Infant, juvenile, subadult, or adult. |
| Aggregate Entropy (AggE) † | Total amount of disorder (unitless). |
| Average Entropy (AvgE) † | Disorder calculated for each time slice, then averaged over the total sound (unitless). |
| Bandwidth 90% (BW) † | (Frequency 95%) – (Frequency 5%) (Hz). |
| Center Frequency (CF) † | Frequency at which the energy within the selection is divided equally in two (Hz). |
| Center Time (CT) † | Time at which the energy within the selection is divided equally in two (seconds). |
| Delta Time (DT) | Time between the start and end points of the selection (seconds). |
| Duration 90% (Dur) † | (Time 95%) – (Time 5%) (seconds). |
| First/Last Syllable (F/L) | Designation of initial and final syllables of the unit, excluding introductory and terminal syllables. |
| Introductory/Terminal Syllable (I/T) | Subunits at the beginning and/or end of a unit that were a) a markedly different shape than the rest of the unit's syllables; and/or b) had start frequencies that did not follow the overall arc of the unit (Robakis et al., 2018). Not present in all units. |
| Frequency 5%, Frequency 95% (F5, F95) † | Frequencies within a selection at 5% and 95% of the energy in the call (Hz). |
| Mass | Body weight taken during mark-recapture (grams). |
| Maximum Frequency (MF) † | Frequency with the maximum amount of energy in the selection (Hz). |
| Maximum Frequency Syllable (MFSyll) | Syllable with the maximum frequency (binary designation). |
| Maximum Power (MP) † | Time of maximum amplitude in the selection (seconds) |
| Maximum Power Syllable (MPSyll) | Syllable with maximum power (binary designation). |
| Subunit Number (Subunit) | Ordinal numbers (1...N) assigned to each discrete syllable in a unit. |
| Time 5%, Time 95% (T5, T95) † | Time of 5% and 95% of the energy in the selection (seconds). |
| Unit Number (Unit) | Unique numerical identifier belonging to each long call. |
| Year | Year (2014-2016) the recording was made. |

Table S1 (continued)
| **Unit Set** |  |
| --- | --- |
| ***Variable*** | ***Description*** |
| Age Class | Infant, juvenile, subadult, adult. |
| Delivery Rate (DelRate) | (Number of Subunits)/(Total Duration). |
| Duration Average (DurAv) | Mean of Duration 90% over all syllables. |
| Number of Introductory Syllables (IntroNo) | Total number of introductory syllables in the unit. |
| Intersyllable Interval Average (IntAv) | [(Time 5%) - (Time 95% of preceding syllable)]/(Number of Subunits). |
| Mass | Body weight taken during mark-recapture (grams). |
| Maximum Frequency Syllable Number (MFNo) | Subunit Number which contains the maximum frequency. |
| Maximum Frequency Syllable Index (MFSI) | (Maximum Frequency Syllable)/(Total # of Subunits). |
| Maximum Power Syllable Number (MPNo) | Subunit Number of the subunit which contains the maximum power. |
| Maximum Power Syllable Index (MPSI) | (Maximum Power Syllable)/(Total # of Subunits). |
| Number of Subunits (SubNo) | Total number of discrete syllables within each unit. |
| Subunits Calculated (SubCalc) | (Number of Subunits) – (Introductory and terminal syllables). |
| Number of Terminal Syllables (TermNo) | Total number of terminal syllables within the unit. |
| Total Duration (TD) | (Time 95% of last syllable) – (Time 5% of first syllable). |
| Unit Number (Unit) | Unique numerical identifier belonging to each long call. |
| Year | Year (2014-2016) the recording was made. |

### Appendix 2

Table S2. Ethogram of behavioral responses of emperor tamarins to playback experiments. Behaviors were categorized as either reactive or non-reactive. Behaviors included in the reactive category are responses to disturbances or startling stimuli commonly observed in callitrichids.

| **Response** | **Behavior** | **Description** |
| --- | --- | --- |
| *Reactive* | Scan | Looking around, moving head rapidly in all directions. |
|  | Attention | Moving head to face speaker, may include rapid head movement to look around in the direction of the speaker. |
|  | Approach | Locomotion (including jumping) in the direction of the speaker, on any axis. |
|  | Withdraw | Locomotion (including jumping) away from the speaker, on any axis. |
|  | Scentmark | Rubbing or sliding the suprapubic, sternal, or anal scent gland on a substrate, such as a branch. |
|  | Aggression | Agonism, such as chasing or hitting, directed toward a conspecific or animal(s) of another species. |
| *Non-reactive* | Forage | Searching for, touching, or inspecting a food source (e.g. fruit, saps, insects), or actively chewing and ingesting something. |
|  | Groom | Combing, scratching, or inspecting the body or hair of another individual; includes receiving grooming, and autogrooming. |
|  | Movement | Locomotion (including jumping) neither towards nor away from the speaker, such as lateral movement on a branch parallel to the speaker. |
|  | Rest | Standing still, sitting, or laying down; eyes may be open or closed. |

### Appendix 3

Proportion of total behaviors exhibited by emperor tamarins following playback conditions (long calls of primary or secondary breeding females) that were classified as reactive or non-reactive (ethogram: Appendix 2). SBS = sex-breeding status class; PBF = primary breeding female; PBM = primary breeding male, SBF = secondary breeding female; SBM = secondary breeding male.

| Condition | S-BS | N | Response | Proportion |
| --- | --- | --- | --- | --- |
| **PBF** | PBF | 17 | Non-Reactive | 0.88 |
|  | PBF |  | Reactive | 0.12 |
|  | PBM | 18 | Non-Reactive | 0.72 |
|  | PBM |  | Reactive | 0.28 |
|  | SBF | 8 | Non-Reactive | 0.50 |
|  | SBF |  | Reactive | 0.50 |
|  | SBM | 6 | Non-Reactive | 0.50 |
|  | SBM |  | Reactive | 0.50 |
| **SBF** | PBF | 14 | Non-Reactive | 0.29 |
|  | PBF |  | Reactive | 0.71 |
|  | PBM | 25 | Non-Reactive | 0.28 |
|  | PBM |  | Reactive | 0.72 |
|  | SBF | 12 | Non-Reactive | 0.42 |
|  | SBF |  | Reactive | 0.58 |
|  | SBM | 1 | Non-Reactive | 1.00 |
|  | SBM |  | Reactive | 0.00 |

